# Intestinal ferritinophagy is regulated by HIF-2α and is essential for systemic iron homeostasis

**DOI:** 10.1101/2020.11.01.364059

**Authors:** Nupur K Das, Amanda Sankar, Andrew J Schwartz, Sumeet Solanki, Xiaoya Ma, Sanjana Parimi, Naiara Santana-Codina, Joseph D. Mancias, Yatrik M Shah

## Abstract

Iron is critical for many processes including oxygen transport and erythropoiesis. Transcriptomic analysis demonstrates that HIF-2α regulates over 90% of all transcripts induced following iron deficiency in the intestine. However, beyond divalent metal transporter 1 (DMT1), ferroportin 1 (Fpn1) and duodenal cytochrome b (Dcytb), no other genes/pathways have been critically assessed with respects to their importance in intestinal iron absorption. Ferritinophagy is associated with cargo specific autophagic breakdown of ferritin and subsequent release of iron. We show here that nuclear receptor co-activator 4 (NCOA4)-mediated intestinal ferritinophagy is integrated to systemic iron demand via HIF-2α. Duodenal NCOA4 expression is regulated by HIF-2α during high systemic iron demands. Moreover, overexpression of intestinal HIF-2α is sufficient to activate NCOA4 and promote lysosomal degradation of ferritin. Promoter analysis revealed NCOA4 as a direct HIF-2α target. To demonstrate the importance of intestinal HIF-2α/ferritinophagy axis in systemic iron homeostasis, whole body and intestine-specific NCOA4-null mouse lines were assessed. These analyses demonstrate an iron sequestration in the enterocytes, and significantly high tissue ferritin levels in the dietary iron deficiency and acute hemolytic anemia models. Together, our data suggests efficient ferritinophagy is critical for intestinal iron absorption and systemic iron homeostasis.

## Introduction

Iron is essential for almost all forms of life as it takes part in numerous cellular processes including oxygen transport, ATP production and DNA synthesis (1). Mammalian iron homeostasis is tightly maintained as both its deficiency and excess are detrimental for health. Iron deficiency is the main cause of anemia worldwide. Proper acquisition of dietary iron is critical for maintaining systemic iron balance. At the level of duodenum, iron is transported into the intestinal epithelial cells (enterocytes) by apical transporters divalent metal transporter 1 (DMT1) and duodenal cytochrome ferric reductase (Dcytb) and exported to the systemic circulation by basolateral transporters ferroportin 1 (Fpn1). Intestinal iron homeostasis is essentially regulated by a hypoxia inducible factor (HIF)-2α, an oxygen- and iron-dependent transcription factor (2).

In addition to iron influx and efflux mechanisms, another integral component of intracellular iron homeostasis is the iron storage function, achieved by ferritin (FTN). FTN stores the surplus cytosolic iron and releases it during conditions of increased requirement or deficiency(3). Iron release from FTN is via autophagic degradation in the lysosome by nuclear receptor coactivator 4 (NCOA4), a process termed as ferritinophagy (4,5). Ferritinophagy occupies a critical place in systemic iron homeostasis and is itself regulated by cellular iron levels (6). More recent discoveries about the role of NCOA4 in erythropoiesis highlights its connection with mammalian oxygen carrying function (7-9).

We and others have previously characterized the significance of intestinal handling of dietary iron with respects to systemic regulation and erythropoiesis (10). In addition, the role of intestinal FTN in iron absorption is also well studied (11). But question still remains how the enterocyte ferritinophagic pathway plays any role in systemic iron regulation. Enterocyte membrane iron flux is essentially regulated by master transcription factor HIF-2α. In this work, we first sought for the potential connection between intestinal HIF signaling and ferritinophagic pathways. Using relevant genetic mouse models, that includes constitutive and conditional NCOA4 knockout mice, we then investigated the role of intestinal NCOA4 in the pathophysiology of systemic iron disorders.

## Results

### Intestinal NCOA4 expression is induced in conditions of high systemic iron demand

Dietary iron deficiency and increased erythropoiesis are the two major conditions that create high systemic iron demand. Ferritin breakdown is a major mechanisms by which intracellular iron level is restored (12). To study the role of ferritinophagy during high systemic iron demand, NCOA4 expression was assessed in iron-deficient (< 5 ppm iron diet, 2 weeks; 350 ppm iron diet was used as control) and phenylhydrazine (Phz)-treated mice by q-RTPCR analyses in tissues central to systemic iron homeostasis, e.g. absorption (duodenum), storage (liver) and recycling (spleen). Duodenal NCOA4 expression was significantly increased in both the conditions (**Figure 1A**). Hepatic NOCA4 levels did not show significant change (**Figure 1B**). With regards to splenic expression, iron-deficiency had no significant effect, while Phz-treatment resulted in a significant increase (about 5-7-fold) (**Figure 1C**). Also, mRNA expression analyses of duodenal glycolytic (pyruvate dehydrogenase kinase isozyme 1 (PDK1), and phosphoglycerate kinase 1(PGK1)glucose transporter 1 (Glut1)), and two other HIF target genes, transferrin receptor 1 (TfR1) and ankyrin repeat domain 37 (ANKRD37) confirm induction of HIF activity during both dietary iron deficiency and acute hemolytic anemia (**Figure S1A &B**). Together, these data suggest that, ferritinophagy is highly active at the intestinal level in conditions of both local and systemic iron deficit. High splenic NCOA4 level in Phz-treatment reflects the high local iron turnover during massive hemolysis.

**Figure 1.**
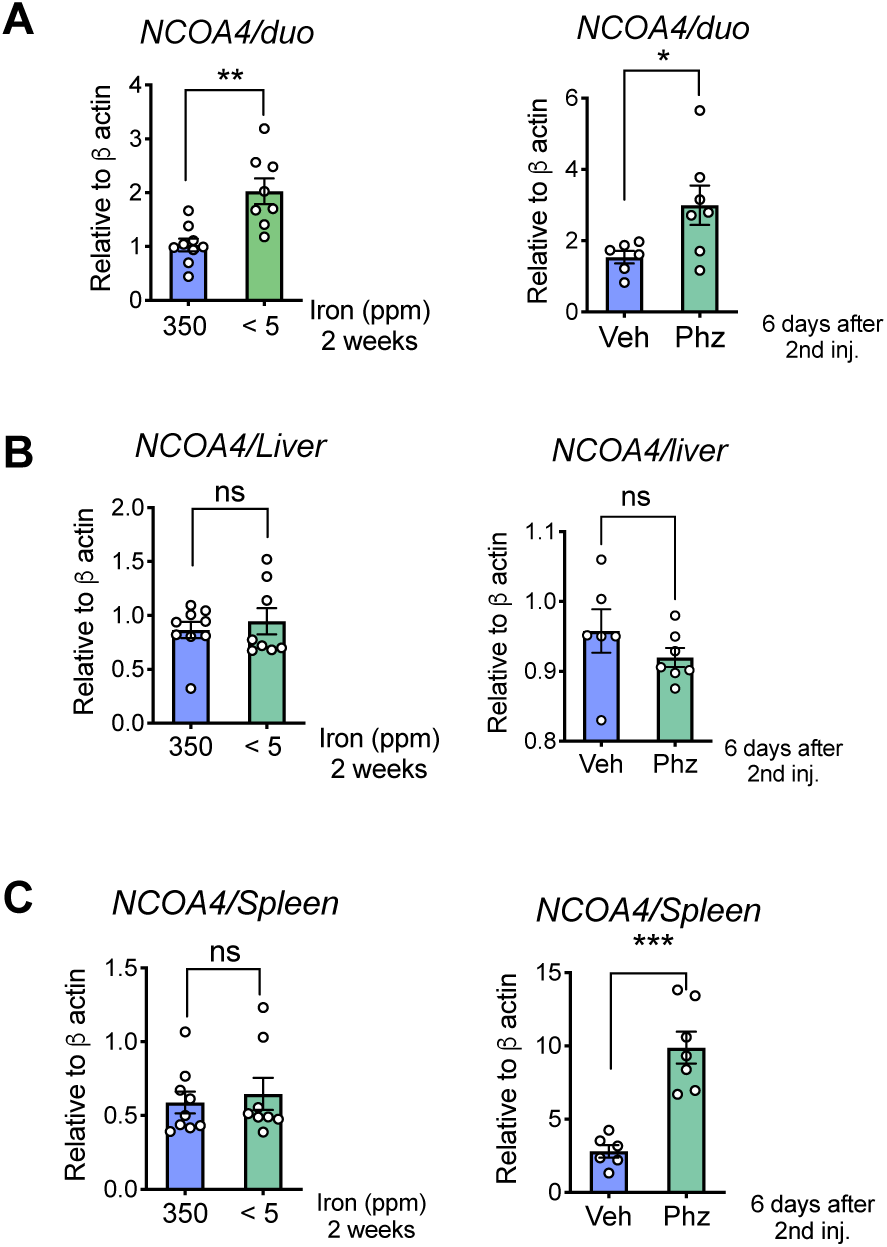
Intestinal *Ncoa4* expression is induced by high systemic iron demand. Mice were treated with iron diets (350 vs. <5 ppm) or phenylhydrazine (Phz) i.p., and NCOA4 gene expression analysis was performed in duodenum (**A**), liver (**B**) and spleen (**C**). All data are mean ± SEM. t test was performed for statistical analysis. ^∗^p < 0.05, ^∗∗^p < 0.01, ^∗∗∗^p < 0.001.

### Intestinal NCOA4 expression is essentially dependent on HIF-2α

A dynamic and coordinated interplay between cellular iron absorption and storage is critical for maintaining systemic iron level. The major pathway by which systemic iron deficit is corrected is via HIF-2α mediated increase of intestinal iron absorption (13). Induction of intestinal NCOA4 expression at the protein level during dietary iron deficiency was confirmed by Western analyses (**Figure 2A**). To study the possible association between HIF-2α and ferritinophagy, NCOA4 expression was assessed in wild type (HIF-2α^*F/F*^) and intestine-specific HIF-2α knockout (HIF-2α^*ΔIE*^) mice fed on 350- and < 5 ppm iron diet for 2 weeks. In agreement with our initial data (**Figure 1A**), duodenal NCOA4 RNA expression was significantly increased in the HIF-2α^*F/F*^ mice, while the effect was abolished in the HIF-2α^*ΔIE*^ cohort (**Figure 2B**). This data is consistent with expression of DMT1, a well-characterized HIF-2α target gene (**Figure 2B**). Western analysis confirmed that induction of duodenal NCOA4 protein level is HIF-2α dependent (**Figure 2C**). Further analysis using both pharmacological (FG4592; prolylhydroxylase (PHD) inhibitor) and genetic (Vhl^*ΔIE*^) models revealed that intestinal NCOA4 RNA expression positively correlates with the generalized HIF induction in the intestine (**Figure 2D**). NCOA4 protein expression in Vhl^*ΔIE*^ duodenum was found to be highly increased compared to that of wild type (Vhl^*F/F*^) tissue (**Figure 2E**). Next, to study the role of NCOA4 in ferritin turnover, NCOA4 and ferritin (FTN) expressions were analyzed in the lysosomal fraction of duodenal lysates. Increased lysosomal FTN expression correlated with the induction of NCOA4 in the Vhl^*ΔIE*^ samples, highlighting the role of NCOA4 in lysosomal enrichment of FTN for subsequent degradation (**Figure 2F**). To further confirm that intestinal expression of NCOA4 is specifically dependent on HIF-2α, intestinal HIF-2α was deleted in the Vhl^*ΔIE*^ mice (Vhl/ HIF-2α^*ΔIE*^). Consistent with the duodenal DMT1 RNA expression, NCOA4 induction was also abolished in the Vhl^*ΔIE*^ mice lacking HIF-2α, suggesting the NCOA4 expression is essentially dependent on HIF-2α (**Figure 2G**). Western analysis confirmed that HIF-2α is essential for intestinal NCOA4 expression (**Figure 2H**). Next, to address whether NCOA4 is a target of HIF-2α, 1kb mouse NCOA4 promoter was cloned into pGL3 luciferase vector and the promoter activity was assessed in either HIF-1α or HIF-2α overexpressed HCT116 human intestinal cell line. Luciferase activity was increased by about 2.5-3-fold in the HIF-2α overexpression, while HIF-1α overexpression had no significant change compared to empty vector control (**Figure 2I**). It is important to note here that the fold increase in the NCOA4 promoter activity is equivalent to its induction in RNA expression during high iron demand (**Figure 1A**). Together, these data establish that intestinal NCOA4 is a novel target of HIF-2α.

**Figure 2.**
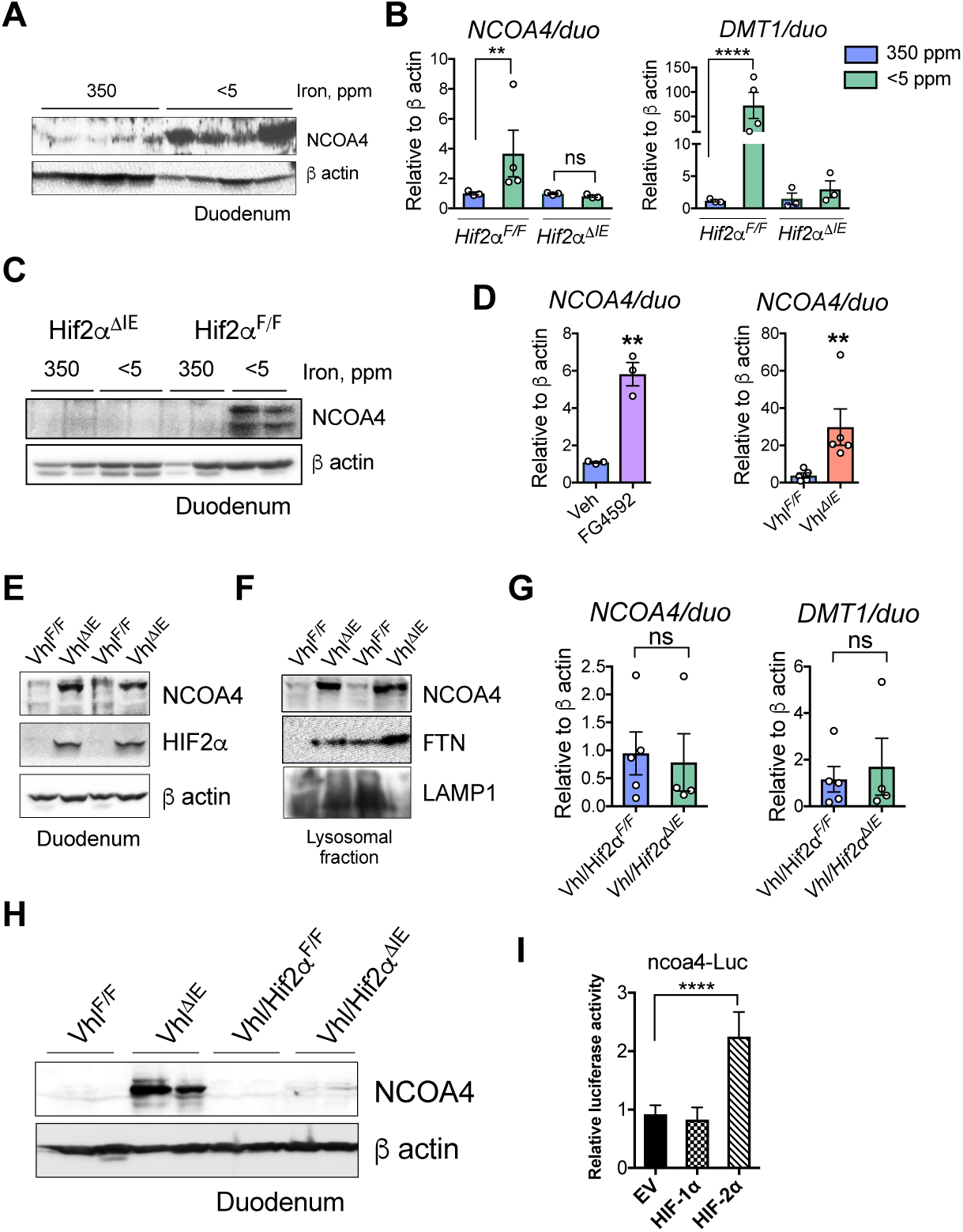
Intestinal NCOA4 expression is HIF2α-dependent. **A**. Wild type mice were treated with 350- or <5 ppm iron diets for 2 weeks followed by NCOA4 Western analysis of duodenum. **B**. Wild type (HIF-2α^*F/F*^) and intestine-specific HIF-2α (HIF-2α^*ΔIE*^) knockout mice were treated with 350- or <5 ppm iron diets for 2 weeks followed by NCOA4 and DMT1 gene expression analysis of duodenum. **C**. Duodenal NCOA4 Western analysis of HIF-2α^*F/F*^ and HIF-2α^*ΔIE*^ mice fed with 350- or <5 ppm iron diets for 2 weeks. **D**. Duodenal NCOA4 gene expression in FG4592-treated mice or intestine-specific VHL knockout (Vhl^*ΔIE*^) mice. **E**. Duodenal NCOA4 and HIF-2α Western analysis in wild type (Vhl^*F/F*^) and Vhl^*ΔIE*^ mice. **F**. Duodenal NCOA4 and FTN Western analysis in Vhl^*F/F*^and Vhl^*ΔIE*^ mice from lysosomal fraction. **G**. Duodenal NCOA4 and DMT1 gene expression analysis in wild type (Vhl/HIF-2α^*F/F*^) and intestine-specific VHL-HIF-2α double knockout (Vhl/ HIF-2α^*ΔIE*^) mice. **H**. Duodenal NCOA4 Western analysis in wild type and Vhl/ HIF-2α^*ΔIE*^ mice. **I**. NCOA4 promoter analysis by luciferase assay in HCT116 cells co-transfected with empty vector (EV), HIF-1α or HIF-2α. All data are mean ± SEM. t test was performed for statistical analysis. ^∗^p < 0.05, ^∗∗^p < 0.01, ^∗∗∗^p < 0.001, ^∗∗∗∗^p < 0.001.

### Systemic NCOA4 deficiency causes tissue iron sequestration and mild microcytosis

Previous reports have shown that whole body NCOA4 deficiency has variable effects on systemic iron parameters (6,8). To characterize and confirm the role of NCOA4 in systemic iron homeostasis, we have generated a NCOA4 knockout (KO) mouse line by using CRISPR-mediated guide RNA (**Figure 3A-C**). Consistent with previous reports, successful deletion of NCOA4 (**Figure 3D and S2A**) resulted in tissue iron sequestration evidenced by increased FTN expression (**Figure 3E**) and enhanced Prussian blue staining (**Figure 3F**) in the NCOA4 KO duodenum, liver and spleen. CBC analysis in NCOA4 KO mice exhibited significantly lower MCV and MCH, compared to those of wild type littermates, suggesting microcytosis and erythrocytosis (**Figure 3G**). Interestingly, the KO mice did not show a significant change in hemoglobin concentration (**Figure 3G**), indicating systemic NOCA4 deficiency does not cause major systemic iron deficit in basal conditions, although it causes appreciable tissue iron sequestration. It is worthwhile to note here that NCOA4 KO mice do not show significant changes in hepatic hepcidin, but significantly lower renal erythropoietin (EPO) expression, compared to the wild type cohort (**Figure S2B**).

**Figure 3.**
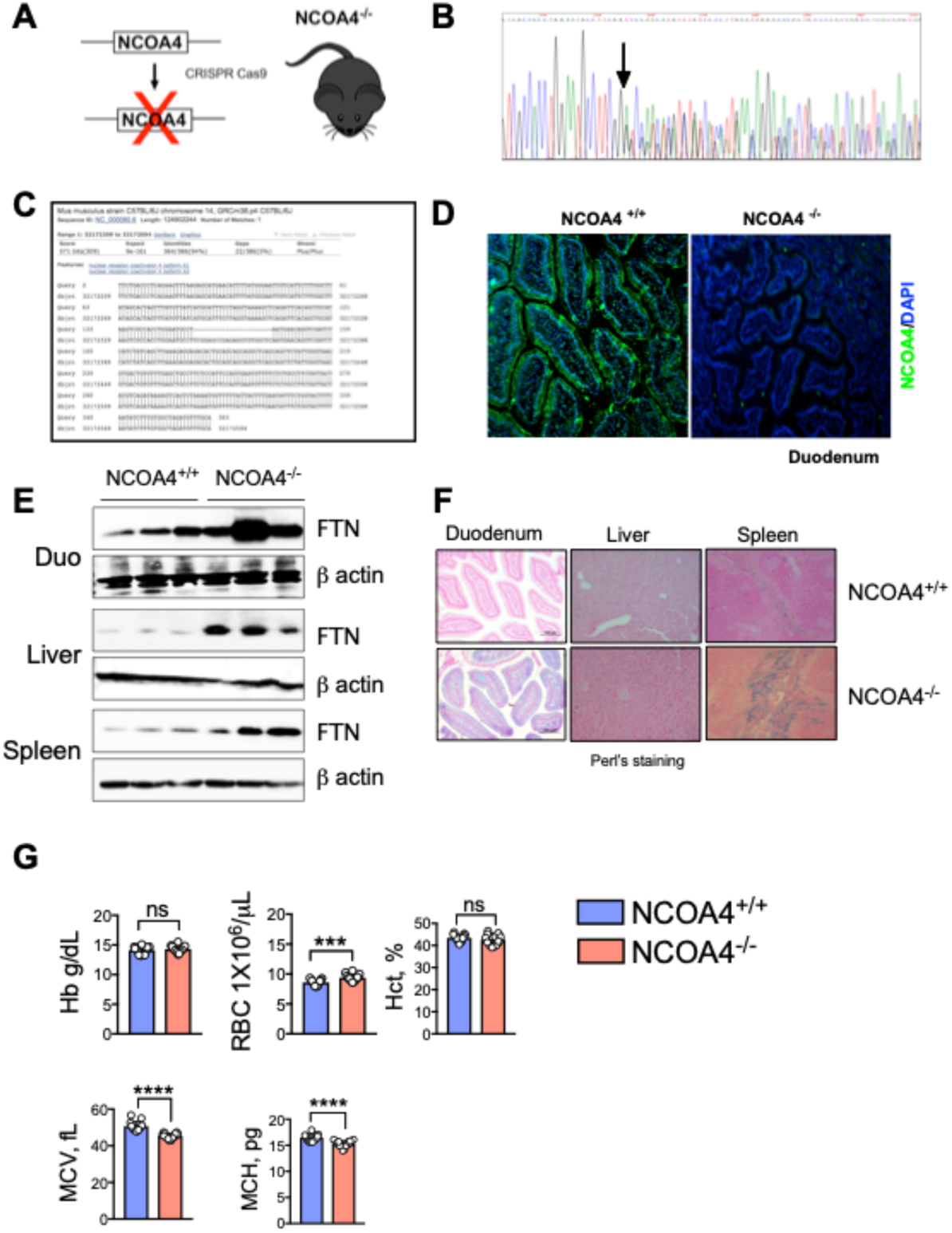
Systemic NCOA4 deficiency causes tissue iron sequestration and mild microcytosis. **A**. Schematic depicting the generation of NCOA4 knockout (KO) mouse line. **B**. DNA sequence chromatogram showing peaks-on-peaks (arrow) of NCOA4 KO mouse. **C**. Representative DNA sequence analysis by NCBI BLAST of NCOA4 KO mouse. **D**. Immunohistochemical analysis of NCOA4 in wild type and NCOA4 KO duodenum. FTN Western analysis (**E**) and Perl’s staining (**F**) in wild type and NCOA4 KO duodenum, liver and spleen. G. Complete blood count (CBC) analysis in wild type and NCOA4 mice. All data are mean ± SEM. t test was performed for statistical analysis. ^∗^p < 0.05, ^∗∗^p < 0.01, ^∗∗∗^p < 0.001, ^∗∗∗∗^p < 0.001.

### Whole-body NCOA4 disruption causes tissue iron sequestration and is required to sustain iron levels in hemolytic anemia, but not in dietary iron deficiency

Since NCOA4 deficiency did not exhibit a major phenotype in basal iron conditions, we inquired whether conditions causing high systemic iron demand could implicate NCOA4. Phz causes massive hemolysis and an acute deficit in systemic iron level caused by increased erythropoiesis. Consistent with this, FTN expression was drastically reduced in the duodenum and liver, but not in the spleen, of Phz-treated wild type mice (**Figure 4A**). Phz-treated NCOA4 KO mice exhibited significantly more tissue FTN expression, suggesting NCOA4 is essential for iron mobilization is during high systemic iron demand (**Figure 4A**). To study how NCOA4 deficiency regulates systemic iron levels following acute increase in iron demand, we performed CBC analysis of a time course study of Phz treatment (0, 3 and 6 days following PhZ treatment). Interestingly, Phz treatment was able to alter the basal differences in CBC parameters (Figures 4B). NCOA4 KO mice exhibit significantly lowered hemoglobin and hematocrit, after 6 days of Phz treatment, highlighting the role of ferritinophagy in acute iron demand. Further analysis of apical iron transporters, namely DMT1 and Dcytb, show that DMT1 expression is significantly induced in the Phz-treated KO mice, compared to the untreated KO cohort (**Figure 4C**).

**Figure 4.**
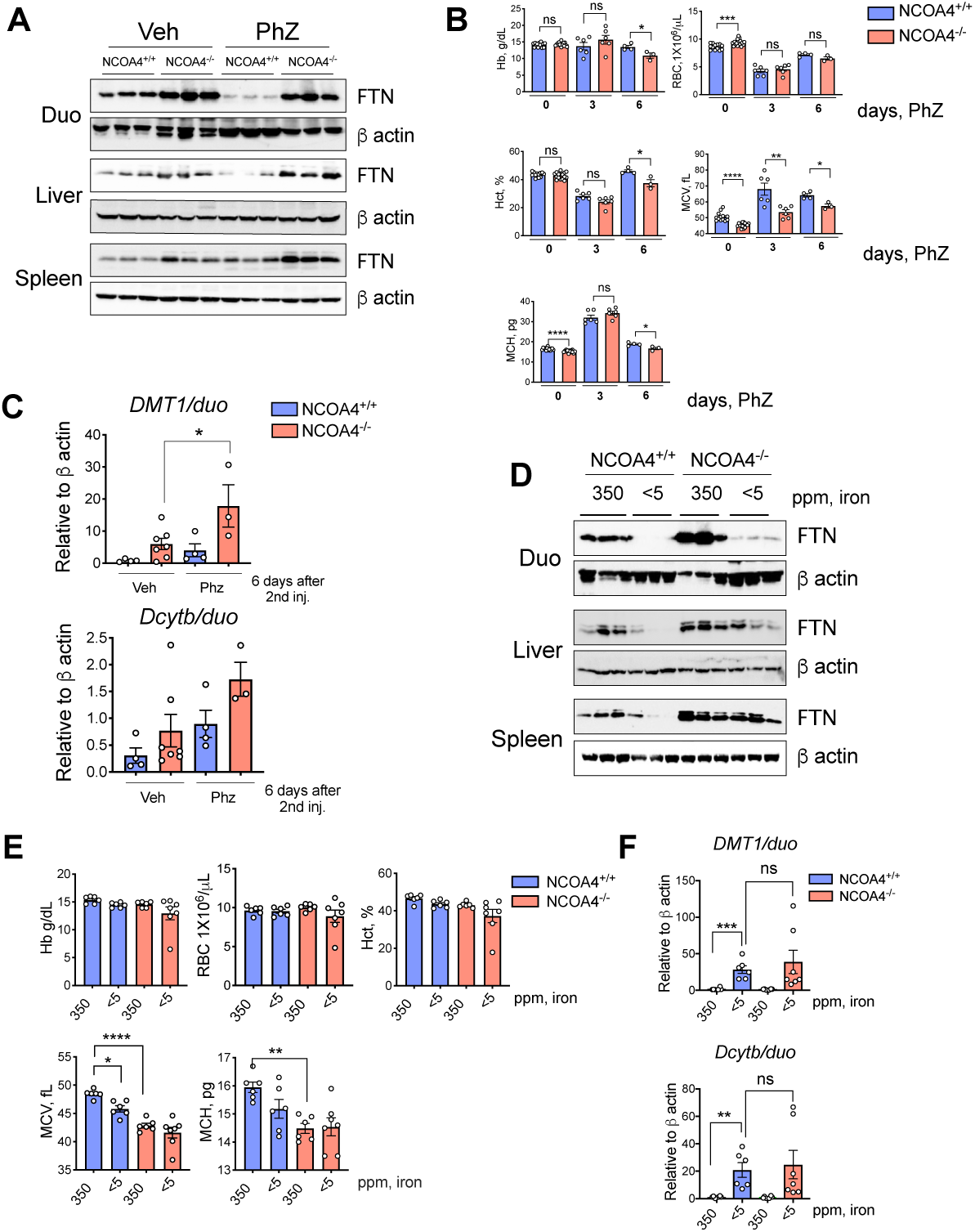
Whole-body NCOA4 disruption alters stress-induced erythropoiesis. Wild type and NCOA4 KO mice were treated with vehicle (Veh) or phenylhydrazine (Phz) followed by FTN Western analysis in duodenum, liver and spleen (**A**); time-dependent CBC analysis (**B**); and duodenal DMT1 and Dcytb gene expression analysis at indicated time point (**C**). Wild type and NCOA4 KO mice were treated with 350- or <5 ppm iron diet for 5 days followed by FTN Western analysis in duodenum, liver and spleen (**D**); time-dependent CBC analysis (**E**); and duodenal DMT1 and Dcytb gene expression analysis at indicated time point (**F**). All data are mean ± SEM. t test was performed for statistical analysis. ^∗^p < 0.05, ^∗∗^p < 0.01, ^∗∗∗^p < 0.001, ^∗∗∗∗^p < 0.001.

Compared to Phz-induced acute hemolytic anemia, dietary iron deficiency is slower in onset and progression. Iron-deficient diet (< 5 ppm) takes about 2 weeks to induce fully develop anemia in mice, but tissue FTN levels drastically fall within one-week. To assess whether NCOA4 deficiency aggravates dietary iron deficiency, we chose the 1-week time point for treatment. As expected, iron-deficient wild type tissues exhibited almost undetected FTN expression (**Figure 4D**). Duodenum, liver and spleen FTN expression in iron deficient NCOA4 KO mice are significantly higher compared to the iron deficient wild type controls (**Figure 4D**). Intriguingly, duodenal FTN expression is indicative of very mild iron retention in the enterocytes, suggesting other compensatory mechanisms are in play for FTN degradation (**Figure 4D**). Consistent with the duration of iron deficiency (1 week), wild type mice did not become anemic (**Figure 4E**). While the 350-ppm fed NCOA4 KO mice showed microcytosis (low MCV) and hypochromia (low MCH), iron-deficient KO mice did not exhibit major changes (**Figure 4E**). There was no significant change in duodenal DMT1 and Dcytb RNA expressions, in the iron deficient NCOA4 KO mice (**Figure 4F**). These findings suggest that systemic NCOA4 deficiency, although causes significant tissue iron sequestration, does not aggravate systemic iron deficit. To explore whether constitutive NCOA4 deficiency results in some compensatory mechanisms to correct the alterations in systemic iron levels, we used tamoxifen-inducible whole body NCOA4 knockout (UBC^Cre/ERT2^; NCOA4^F/F^) mice in a time-dependent manner of NCOA4 ablation. RNA expressions of apical iron transporters DMT1 and Dcytb showed a significant difference with in 1 week of tamoxifen injection, but the changes in gene expression were normalized following chronic knockout of NCOA4 (**Figure S3**).

### Intestine specific NCOA4 deficiency minimally alters systemic iron homeostasis

Intestinal ferritin is essential for efficient iron absorption (11). Our data suggest that compared to other tissues, intestinal ferritin is differentially regulated during high systemic iron demand. To explore the role of intestinal ferritinophagy in systemic iron homeostasis, we used tamoxifen-inducible intestine-specific NCOA4 KO (NCOA4^*ΔIE*^) mice (**Figure 5A**). NCOA4^*ΔIE*^ duodenum showed increased FTN expression (**Figure 5B**), but the CBC analysis did not show any significant difference compared to wild type mice (**Figure 5C**). Following Phz treatment, NCOA4^*ΔIE*^ mice did not show major changes in CBC parameters except a significant increase in MCV compared to Phz-treated wild type mice (**Figure 5D**). There was no significant difference in DMT1 and Dcytb RNA expression in the Phz-treated NCOA4^*ΔIE*^ mice (**Figure 5E**).

**Figure 5.**
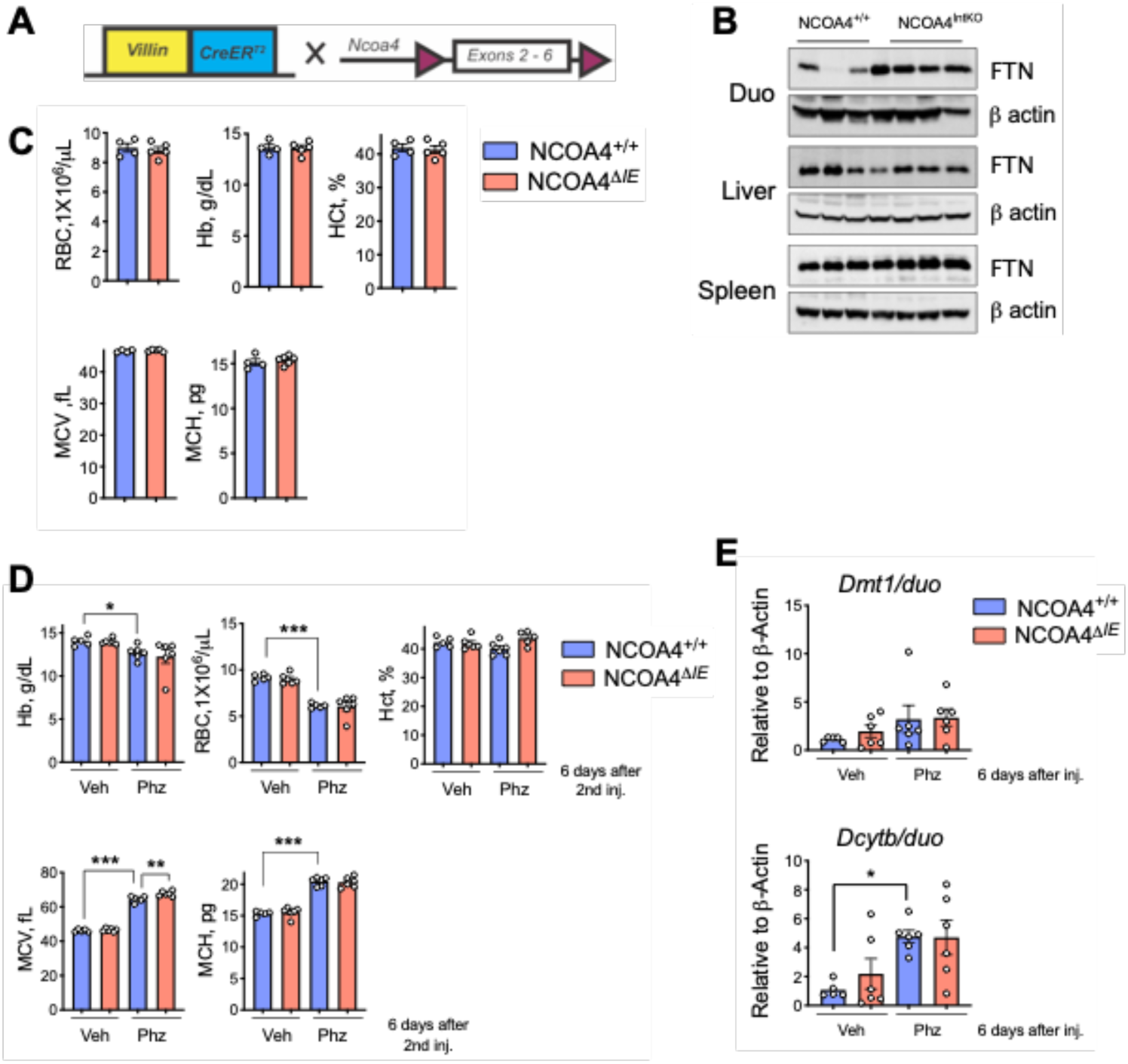
Intestine specific NCOA4 deficiency does not alter systemic iron deficiency. **A**. Schematic depicting the generation of intestine-specific NCOA4 KO mouse line. FTN Western analysis in duodenum, liver and spleen (**B**), and CBC analysis (**C**) of wild type and NCOA4^*ΔIE*^ mice. Wild type and NCOA4^*ΔIE*^ mice were treated with vehicle (Veh) or phenylhydrazine (Phz) followed by CBC analysis (**D**); and duodenal DMT1 and Dcytb gene expression analysis at indicated time point (**E**). All data are mean ± SEM. t test was performed for statistical analysis. ^∗^p < 0.05, ^∗∗^p < 0.01, ^∗∗∗^p < 0.001.

### NCOA4 expression is upregulated during iron hyperabsorption

Analysis of enterocyte, liver and serum iron levels in the NCOA4^*ΔIE*^ mice revealed that significant iron retention takes place in the duodenum (**Figure 6A**). This suggest that NCOA4 is important for efficient mobilization of intestinal iron to the systemic circulation. To assess the potential role of ferritinophagy in iron mobilization, we used liver-specific tamoxifen-inducible hepcidin knockout (HAMP^*ΔLiv*^) mice. We have previously shown that these mice develop rapidly tissue iron overload due to HIF-2α-dependent intestinal hyperabsorption (14). Increase in NCOA4 expression in the duodenum and liver of hepcidin deficient mice suggest that intestinal ferritinophagy is highly active in conditions of hyperabsorption which presumably plays role in efficient iron mobilization to the systemic circulation (**Figure 6B**).

**Figure 6.**
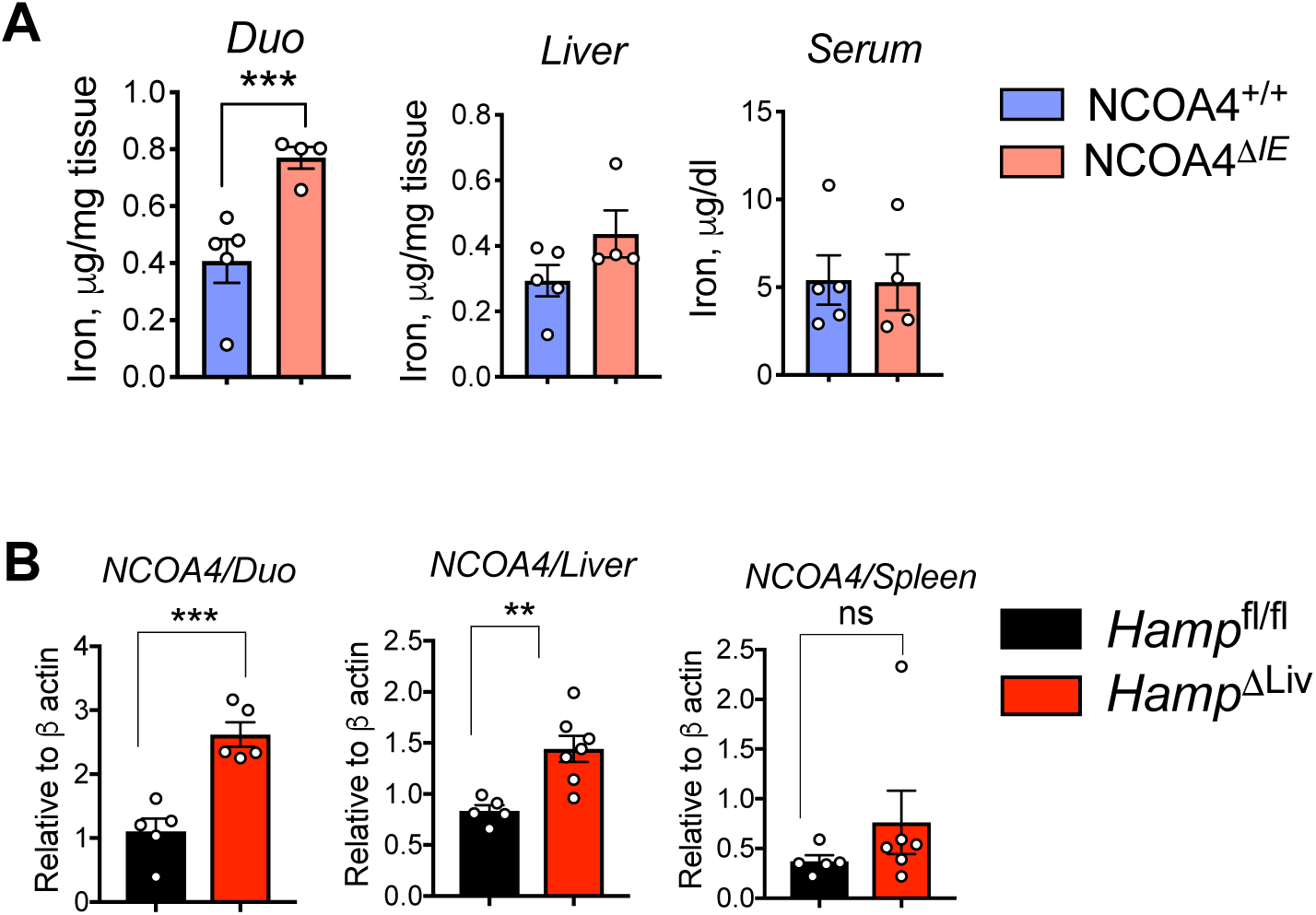
Intestinal NCOA4 is important for enterocyte iron flux. **A**. Tissue (duodenum and liver) and serum iron analysis in wild type vs. NCOA4^ΔIE^ mice. **B**. NCOA4 gene expression analysis in wild type vs. Hamp^ΔLiv^ duodenum, liver and spleen. All data are mean ± SEM. t test was performed for statistical analysis. ^∗^p < 0.05, ^∗∗^p < 0.01, ^∗∗∗^p < 0.001.

## Discussion

Previous studies, using various genetic models of systemic and tissue specific NCOA4 deficiency, have established that NCOA4 is essential for cellular and systemic iron homeostasis. NCOA4 is critical of efficient ferritinophagy *in vivo*, especially during conditions of high iron demand, such as dietary iron deficiency or hemolytic anemia (6,8). A well-orchestrated absorption and intra-enterocyte processing of iron is critical for maintenance of systemic iron parameters (15). In the past decade, the role of HIF-2α in intestinal and systemic iron homeostasis was well defined (13). It was not clear how and whether intestinal iron absorptive functions coordinate with the enterocyte ferritinophagy. In this study, by combining systemic iron deficiency with several genetic models we explored the role of intestinal ferritinophagy in systemic iron regulation.

First, we demonstrate that duodenal NCOA4 expression is consistently induced during high systemic iron demand, such as dietary iron deficiency and acute hemolysis. We and others previously showed that both these conditions are characterized by upregulated HIF-2α function leading to enhanced enterocyte iron flux (2,16). Interestingly, duodenal NCOA4 RNA and protein expression were robustly induced in pharmacological (FG4592, a PHD inhibitor) and genetic HIF overexpression (Vhl^*ΔIE*^) models and were completely abolished in intestine-specific Vhl-HIF2 double KO mice. Subsequent NCOA4 promoter activity analysis in human intestinal cell lines established that intestinal NCOA4 is a specifically dependent on HIF-2α. In contrast to a recent report, where it was demonstrated that hepatic NCOA4 is a target of both HIF-1α and HIF-2α, we show that intestinal NCOA4 expression is specifically dependent on HIF-2α, but not on HIF-1α (17). This exemplifies that ferritinophagy is differentially regulated across different tissues.

To investigate the role of NCOA4 in systemic iron homeostasis, we made whole body NCOA4-KO mouse line utilizing CRISPR-dependent guide RNA for NCOA4. In agreement with previous reports, CBC analysis of our NCOA4-KO line also basally exhibited mild hypochromia with microcytosis. Further stressing the NCOA4-KO mice with Phz-induced hemolysis showed that ferritinophagy was critical maintaining proper erythropoiesis. Interestingly after low iron-diet CBC parameters were not different in wild type and KO mice suggesting the presence of other compensatory mechanisms. It was evident that during systemic iron deficiency, intestinal FTN, but not that of liver or spleen, is mobilized even during absence of NCOA4. This suggested that intestinal ferritin turnover is not entirely dependent on NCOA4.

Enterocyte iron handling is critical for erythropoiesis. Our initial data establishes that high systemic iron demand augments intestinal NCOA4 function in a HIF-2α dependent manner. Erythroid-specific NCOA4 deficiency alters erythropoiesis (8). This work also demonstrated erythroid cell non-autonomous role of NCOA4 in RBC production (8). Based on these observations and our own findings, we hypothesized that intestinal ferritinophagy plays essential role in systemic iron homeostasis.

To investigate the role of NCOA4 in intestinal and as well as systemic iron homeostasis, we then employed an intestine-specific NCOA4 KO mouse model. Basal hematologic parameters of the line did not differ from the whole-body NCOA4-KO mouse model. Stressing them with dietary iron deficiency or acute hemolysis resulted in minimal changes with respect to erythropoiesis and expression of iron transporters.

While we have established that intestinal NCOA4 is a HIF-2α-specific target gene, and its expression is augmented during systemic iron deficiency (where HIF-2α activity is robustly induced), the essentiality of intestinal NCOA4 in these conditions was still uncertain. We utilized liver-specific hepcidin KO mice to focus on the opposite end of the spectrum of high systemic iron demand i.e. systemic iron overload, which is characterized by HIF-2α-mediated hyperabsorption (14). In addition to being dependent on HIF-2α, majority of the players of enterocyte iron metabolism, namely, DMT1, FTN and Fpn1are regulated by iron regulatory protein (IRP) 1 and IRP2 (18). NCOA4 lacks canonical iron response elements (IREs) at the 5’- or 3’ untranslated regions (UTRs) making it unlikely to be regulated by IRE-IRP pathway. Although IRE-IRP mediated FTN regulation and its role in intracellular iron homeostasis is very well characterized (18), it is not clear whether and how this pathway regulates the release of storage iron to the intracellular labile iron pool (LIP).

Together, our data demonstrate that intestinal NCOA4 is a novel target of HIF-2α which plays crucial, although modest, roles in intestinal as well as systemic iron homeostasis. Intestinal iron flux is a function of HIF-2α, and NCOA4 activity is augmented in high HIF-2α activity regardless of the systemic iron levels, strongly indicating intestinal ferritinophagy sits at a critical junction. Further exploration is required to delineate the function of intestinal NCOA4 in the globally prevalent iron disorders.

## Methods

### Cell Culture

HCT116 cell lines were obtained from the American Type Culture Collection company and were grown in Dulbecco’s modified Eagle medium (DMEM) supplemented with 10% fetal bovine serum and 1% anti–anti (Gibco).

### Plasmids

A 1Kb (relative to the transcription start site) mouse NCOA4 promoter region was amplified by PCR from genomic DNA using primers 5’-ATA CAT <u>GCT AGC</u> GCT CTC TAG ACC TCA CGC AG-3’ (forward; underline denotes NheI) and 5’-ATA CAT <u>CTC GAG</u> CCA GAC ACT TAG CCG TGG AA-3’ (reverse; underline denotes XhoI), and was cloned in pGL3 basic vector (Promega). Oxygen stable HIF-1α, HIF-2α overexpression plasmids were described previously (19).

### Animals and treatments

All mice used in this study were on a C57BL/6J background. All other mice were housed in a specific-pathogen free (SPF) facility on a 12-hour light/dark cycle and fed ad libitum with standard chow diet containing ≅ 300 ppm iron. Whole body NCOA4^-/-^ mice, intestine-specific HIF-2α knockout (Hif2α^ΔIE^), intestine-specific VHL knockout (Vhl^ΔIE^), intestine-specific HIF2α-VHL double knockout (Vhl/ Hif2α^ΔIE^) and tamoxifen-inducible liver-specific hepcidin knockout (HAMP^ΔLiv^ : Alb^CreERT2^ ;HAMP^fl/fl^) mice were described previously (14,19).

Tamoxifen-inducible whole body NCOA4 knockout (UBC^Cre/ERT2^; NCOA4^F/F^) mouse lines were provided by Mancias Laboratory. Intestine-specific NCOA4 knockout (NCOA4^ΔIE^ : Vil^CreERT2^; NCOA4^F/F^) mouse line was generated by crossing NCOA4^F/F^ (received from Mancias Laboratory, Boston, MA) with villin-Cre-ER^T2^ mice of the same background (8). To activate the CreER^T2^, mice were intraperitoneal injected with 100 mg/kg of tamoxifen (Sigma) for 5 consecutive days. For iron studies, mice were fed with iron enriched (350 ppm iron), or low-iron (< 5 ppm iron) diet. For hemolysis, mice were injected at 60 mg/kg of body weight on 2 consecutive days and were sacrificed on indicated days following the second injection. FG-4592 or Roxadustat (Cayman chemicals) were administered vial oral gavage at 12.5 mg/kg/day for 5 days. tamoxifen-inducible liver-specific hepcidin knockout (HAMP^ΔLiv^ : Alb^CreERT2^ ;HAMP^fl/fl^)

### Hematological analysis

CBC (complete blood count) analysis was performed by the Unit for Laboratory Animal Medicine Pathology Core at The University of Michigan. Serum and tissue nonheme iron was quantified as described previously (20).

### Iron Staining

Tissue iron detection was performed in formalin-fixed paraffin-embedded sections stained with Perls’ Prussian blue as described previously (20).

### Real-Time quantitative PCR

1μg of total RNA extracted using Trizol reagent from mouse tissues (duodenal epithelial scrapes, liver and spleen), were reverse transcribed to cDNA using SuperScriptTM III First-Strand Synthesis System (Invitrogen). Quantitative PCR (qPCR) reactions were set up in three technical replicates for each sample by combining equal concentration of cDNA, gene-specific forward and reverse primers, with SYBR green master mix, and run in Quant Studio 5 Real-Time PCR System (Applied BioSystems). The fold-change of the genes were calculated using the *ΔΔ*Ct method using beta-actin as the housekeeping gene. The primers are listed in Table S1.

### Western Blot analysis

Cell lysates were collected and lysed in radioimmunoprecipitation assay buffer (50 mM Tris-HCl, 150 mM NaCl, 2 mM ethylenediaminetetraacetic acid, 1% NP-40, and 0.1% TritonX) and protease inhibitors. Protein concentration was estimated through the use of a Bradford protein-dye assay and samples were run in a 10% acrylamide gel. Following transfer to nitrocellulose membrane, the blots were incubated with the following antibodies: NCOA4 antibodies: A302-272A rabbit antibody at 1:1000 (Bethyl Laboratories); sc-30968 goat antibody at 1:200 (Santa Cruz Biotechnology); ferritin rabbit antibody at 1:1000 (Cell Signaling Technology); HIF-2α rabbit antibody (Bethyl laboratories) at 1:1000; actin (Proteintech) at 1:10000; and LAMP1 (Cell Signaling Technology) at 1:1000 dilutions.

### Immunohistochemistry

Duodenums were Swiss-rolled, frozen in cryo-embedding medium, and sectioned at 7 μm. The sections were fixed in 4% paraformaldehyde in PBS and incubated overnight at 4 °C with rabbit anti-NCOA4 antibody (1:100; Bethyl Laboratories. A302-272A) diluted in PBS with 2% BSA. Slides were washed twice with PBS and then incubated for 1 h at room temperature with Alexa Fluor® 488-labeled goat anti-rabbit IgG (1:500; Molecular Probes, Inc., Eugene, OR) diluted in PBS with 2% BSA. After incubation with the secondary antibody, all sections were washed three times for 5 min with PBS and mounted with ProLong® gold antifade reagent with DAPI (Molecular Probes, Inc.). Immunofluorescence was visualized using a Nikon Eclipse TE200 microscope with a ×20 objective. Images were acquired using an Olympus DP71 microscope digital camera and processed using an Olympus DP Controller Version 3.2.1.276 software package (Olympus America Inc., Center Valley, PA).

### Luciferase Assay

HCT116 cells were seeded into a 24-well plate at a cell density of 5 ×10^4^ cells per well. The mouse NCOA4 promoter luciferase construct was co-transfected with oxygen stable HIF-1α, HIF-2α, or empty vector (EV) into cells with polyethylenimine. Cells were lysed in reporter lysis buffer, and firefly luciferase activity was measured using SynergyTM 2 Multi-Mode Microplate Reader (BioTek) and normalized to β-galactosidase (β-gal) activity 48 hours after transfection.

### Statistics

Results are expressed as the mean ± SEM. Significance between 2 groups was tested using a 2-tailed, unpaired *t* test. GraphPad Prism 7.0 was used to conduct the statistical analyses. Statistical significance is described in the figure legends as: ^∗^ p < 0.05, ^∗∗^ p < 0.01, ^∗∗∗^ p < 0.001, ^∗∗∗∗^ p < 0.0001

---

**Figure S1.**
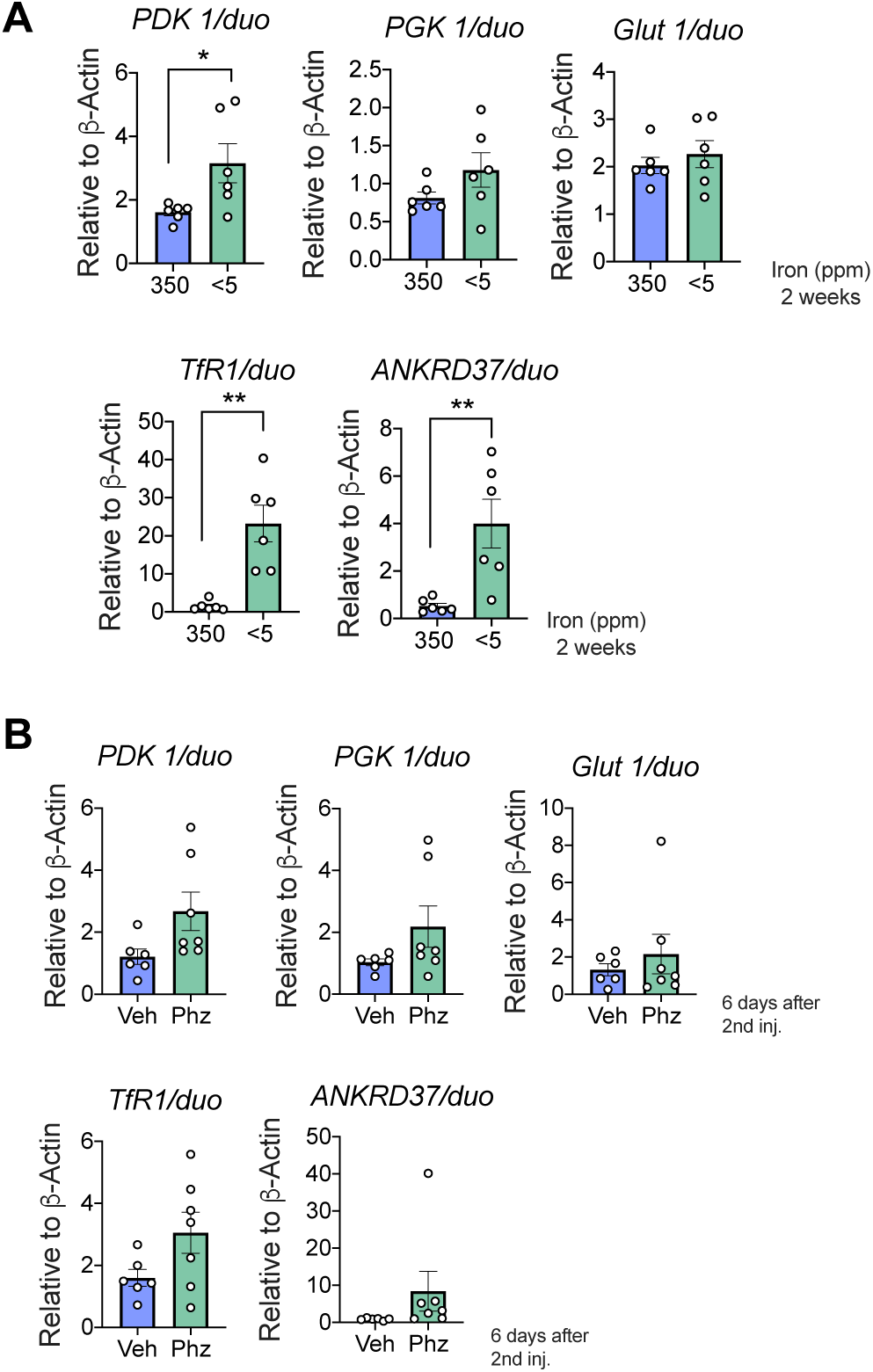
Intestinal HIF activity is induced by high systemic iron demand. Gene expression analysis of duodenal PDK1, PGK1, GLUT1, TfR1 and ANKRD37 in mice treated with iron diets (350 vs. <5 ppm) (**A**), or phenylhydrazine (Phz) i.p. (**B**). All data are mean ± SEM. t test was performed for statistical analysis. ^∗^p < 0.05, ^∗∗^p < 0.01.

**Figure S2.**
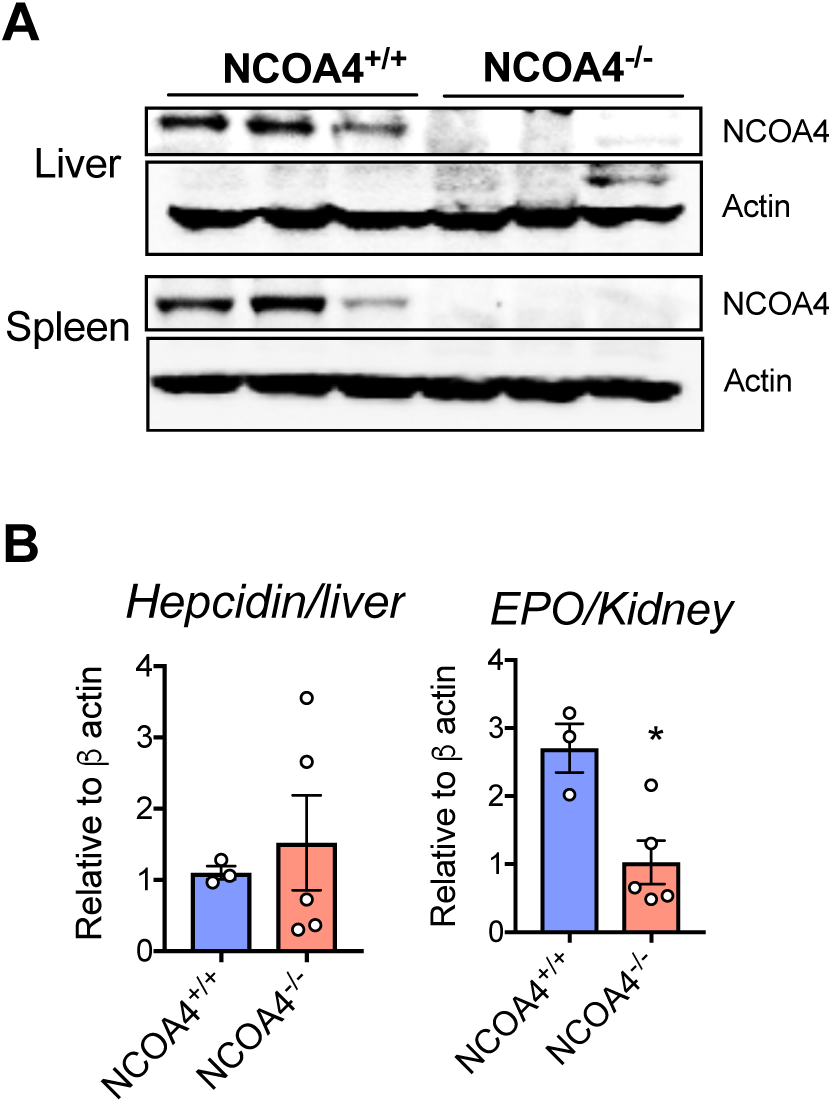
**A**. NCOA4 western analysis in liver and spleen of NCOA4^+/+^ and NCOA4^-/-^ mice. **B**. Gene expression analysis of hepcidin and erythropoietin (EPO) in the liver and kidney respectively, of NCOA4^+/+^ and NCOA4^-/-^ mice. All data are mean ± SEM. t test was performed for statistical analysis. ^∗^p < 0.05.

**Figure S3.**
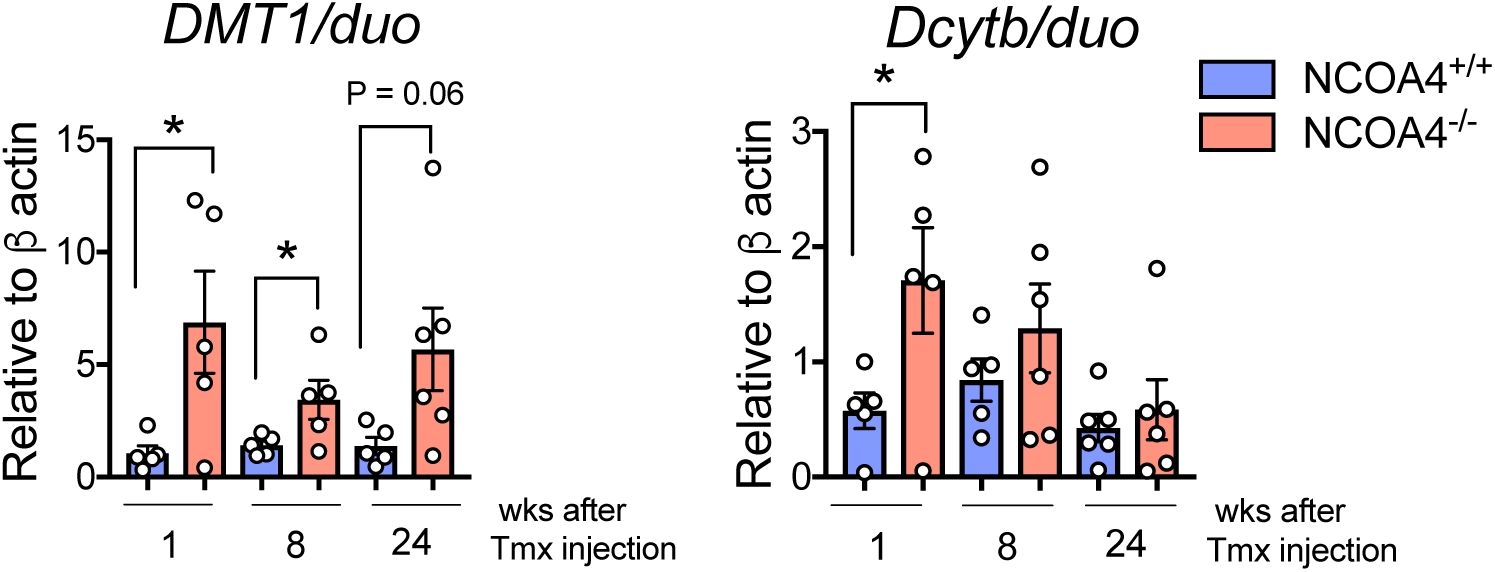
Gene expression analysis of DMT1 and Dcytb in the duodenum, of wild type (NCOA4^+/+^) and tamoxifen-inducible whole body NCOA4 knockout (NCOA4^-/-^ : UBC^Cre/ERT2^; NCOA4^F/F^) mice. All data are mean ± SEM. t test was performed for statistical analysis. ^∗^p < 0.05.

**Table S1.**
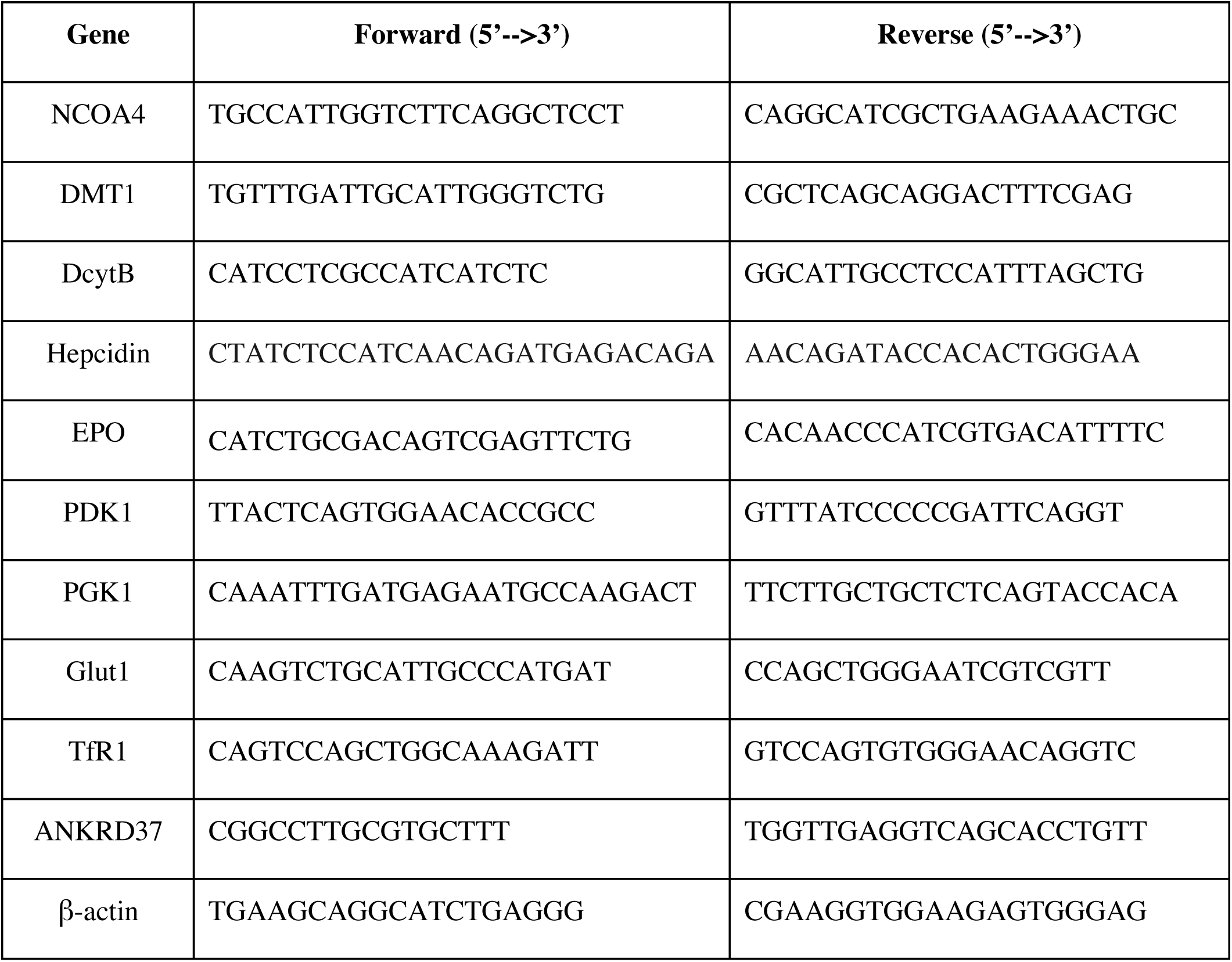
qPCR Primer sequences.

